# Phage proofing *Pseudomonas putida* uncovers novel broad-spectrum phage resistance protein Psh

**DOI:** 10.64898/2026.09.02.748959

**Authors:** Ben Diaz, Stephen Won, James D. Jaryenneh, Md Maksudur Rahman, Vallari R. Chourasia, Catherine M. Mageeney

## Abstract

Broad-spectrum phage resistance offers an important layer of protection against bacterial fermenter crashes during biomanufacturing processes, yet the underlying mechanisms are often poorly defined or come at a fitness cost. Using experimental evolution, we generated two *Pseudomonas putida* strains that were resistant to at least six phage genera. Genome sequencing revealed a single frameshift deletion in each strain that restored functionality of a Type I secretion system (T1SS) ATPase. We also found strong transcriptional upregulation of a nearby protein, PP_1794, which we coin **P**hage **sh**ielding **h**elix rich protein (Psh). Overexpression experiments show that Psh protein is secreted by this restored T1SS to confer complete phage resistance. When compared to wild-type *P. putida* KT2440, no growth or expression defects were evident, but pyoverdine production was reduced. The Psh gene and T1SS are widely distributed across Gram-negative bacteria. These findings uncover a previously uncharacterized phage defense system in *P. putida* based on secretion-mediated receptor masking.

## Introduction

Bacteriophages (phages) are the most abundant biological entity on Earth and infect bacteria by injecting their DNA into a bacterial host to co-opt cell processes and produce progeny phages before lysing the cell. Though phages have useful applications in treating bacterial infections in mammalian and plant species^1^, they can also contaminate bacterial fermentation processes^2–4^, causing collapse of bacterial populations and increased biomanufacturing costs^4,5^. Current solutions to overcome phage-mediated fermenter crashes include extended sanitation, raw material treatments, and improved fermenter design^6,7^. Some modern strategies also focus on evolving or engineering phage-resistant strains^2,3^. Despite these precautions, phages are diverse and abundant, and therefore, continue to represent a potential mode of failure.

Bacteria have access to numerous sophisticated mechanisms to prevent phage infection at each stage of the phage lifecycle. These phage defense systems can block phage adsorption or entry (i.e. receptor modifications), degrade phage nucleic acids (i.e. restriction modification systems and CRISPR Cas systems), target later stages of the phage lifecycle (i.e. tail assemble blocker and capsid remodeling), or trigger cell apoptosis to save the population (i.e. ABI and toxin-anti-toxin)^8,9^. There is no single phage defense that effectively protects against all phages because phages are diverse and can rapidly evolve to overcomes these mechanisms^10^. While these mechanisms effectively prevent phage infection, the associated fitness trade-offs have not been elucidated for most systems. Further, many of these defense systems are clustered together in bacterial genomes which may imply certain combinations are necessary for function or impart fitness tradeoffs^11,12^.

Here, we aimed to develop a phage-resistant *Pseudomonas putida* strain. *P. putida* strains are a common biomanufacturing chassis due to ability to tolerate solvents and organics, and are able to grow on a wide variety of carbon sources, including those important to the bioeconomy such as lignin hydrolysates^13,14^. Additionally, *P. putida* has a well-characterized genome and numerous synthetic biology toolkits that have been used to engineer it to produce diverse products including alcohols, proteins, and polymers^13^. While phages capable of infecting *P. putida* have only recently been isolated^15–17^, many of these phages can disrupt cultures within hours of infection, presenting a significant risk to biomanufacturing processes dependent on this strain.

In this study, *P. putida* KT2440 and mt-2 were challenged with four diverse phage isolates and a three-phage cocktail to facilitate evolved phage resistance. A broad panel of *P. putida* phages were tested against the evolved strains. We discovered that two strains, one KT2440 and one mt-2, were completely resistant to every phage tested with no impact to overall fitness. We used genomic and transcriptomic analysis to aid in understanding the mechanism responsible for this broad phage resistance and uncovered a novel phage defense mechanism. This system consists of a previously uncharacterized protein, PP_1794, that is excreted by a Type I secretion system (T1SS). The PP_1794 gene has an early stop codon in wild-type *P. putida* KT2440 but this open reading frame is restored through single base pair frameshift deletion in our phage-resistant mutants (**Figure 1**). Based on the observed phenotype and its predicted 3D structure, we named PP_1794 **P**hage **sh**ielding **h**elix rich protein (Psh).

**Figure 1.**
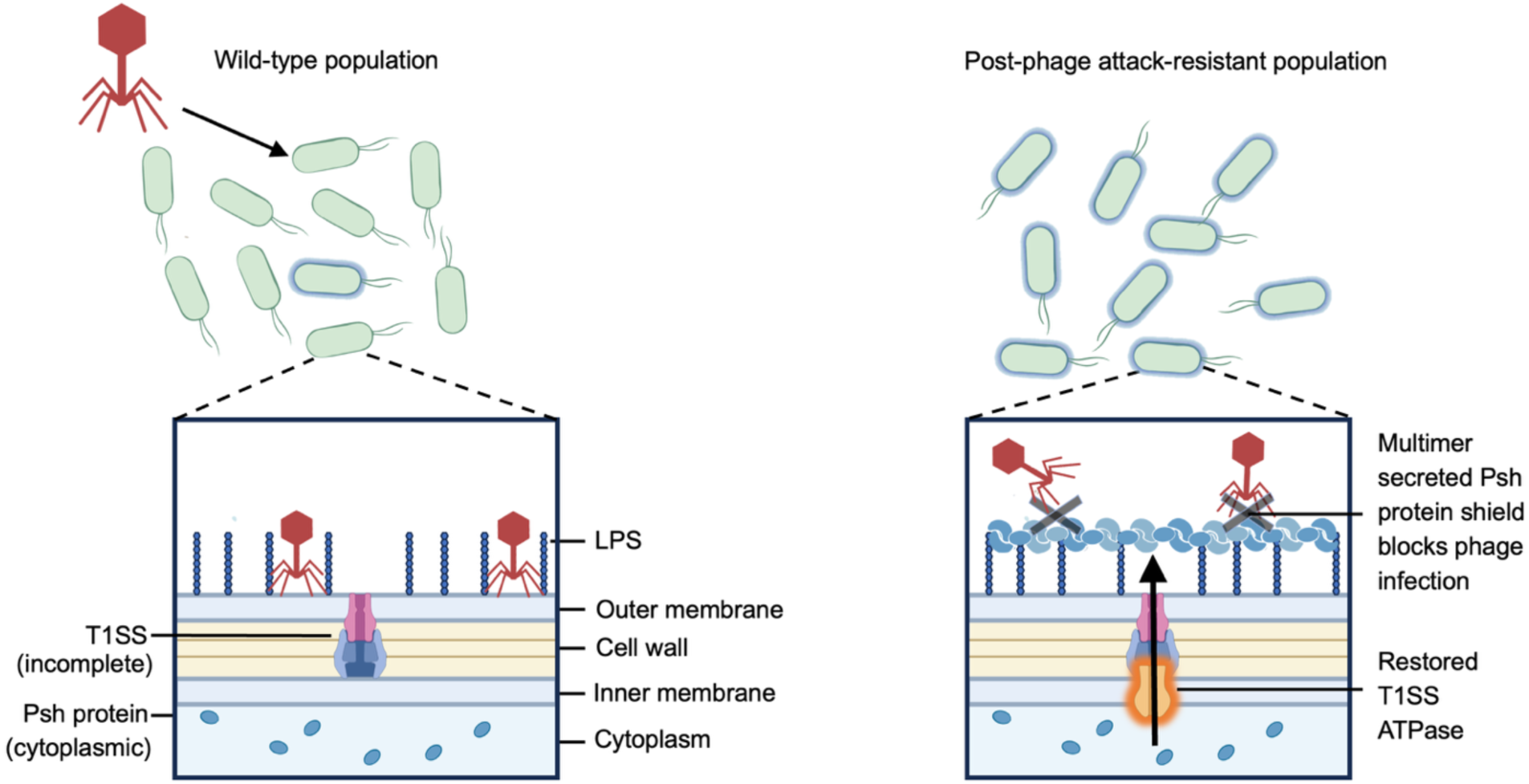
Conceptual model of Psh protein and restored T1SS phage defense mechanism. The bacterial population on the left represents wild-type *P. putida* KT2440, which are phage susceptible, and have an incomplete T1SS. Wild-type KT2400 produces Psh protein but lacks a functional T1SS for export and therefore cannot bind the O-antigen to block phage infection. In the right panel, we show the proposed molecular mechanism for EmajogiR.1. In these resistant bacteria, the *aprD* ATPase of the T1SS is restored by a frameshift deletion, and the Psh protein is exported at high efficiency and forms multimers by binding to the O-antigen creating a steric hinderance for phage infection.

## Results

### Establishing a phage-resistant *P. putida* strain

We employed a two-phase experimental evolution approach to obtain a stable, phage-resistant *P. putida* strain that was still suitable for biomanufacturing. Phages were added daily to a freshly diluted *P. putida*. Each day samples were taken and tested for phage-resistance. If no resistance was observed, we repeated this process using the previous culture until resistance was observed (ranging from 4-7 days) (**Figure 2a**). Any culture that was determined to be resistant with an efficiency of plating (EOP) of more than four, was streaked without the addition of phages to remove residual phages from the culture to create a stable strain. We then tested the stable evolved strains against a diverse panel of phages ^15,16^ and observed two strains resistant to all phages tested (EmajogiR.1 and PLiverwasterR.1) and two strains exhibiting reduced EOP to several phages (MiCathR.1 and MiCathR.2) (**Figure 2b**). We also confirmed that *P. putida* KT2440_EmajogiR.1 (hereafter referred to as EmajogiR.1) was resistant to all phages tested, including a sixteen-phage cocktail in liquid culture (**Figure 2c**).

**Figure 2.**
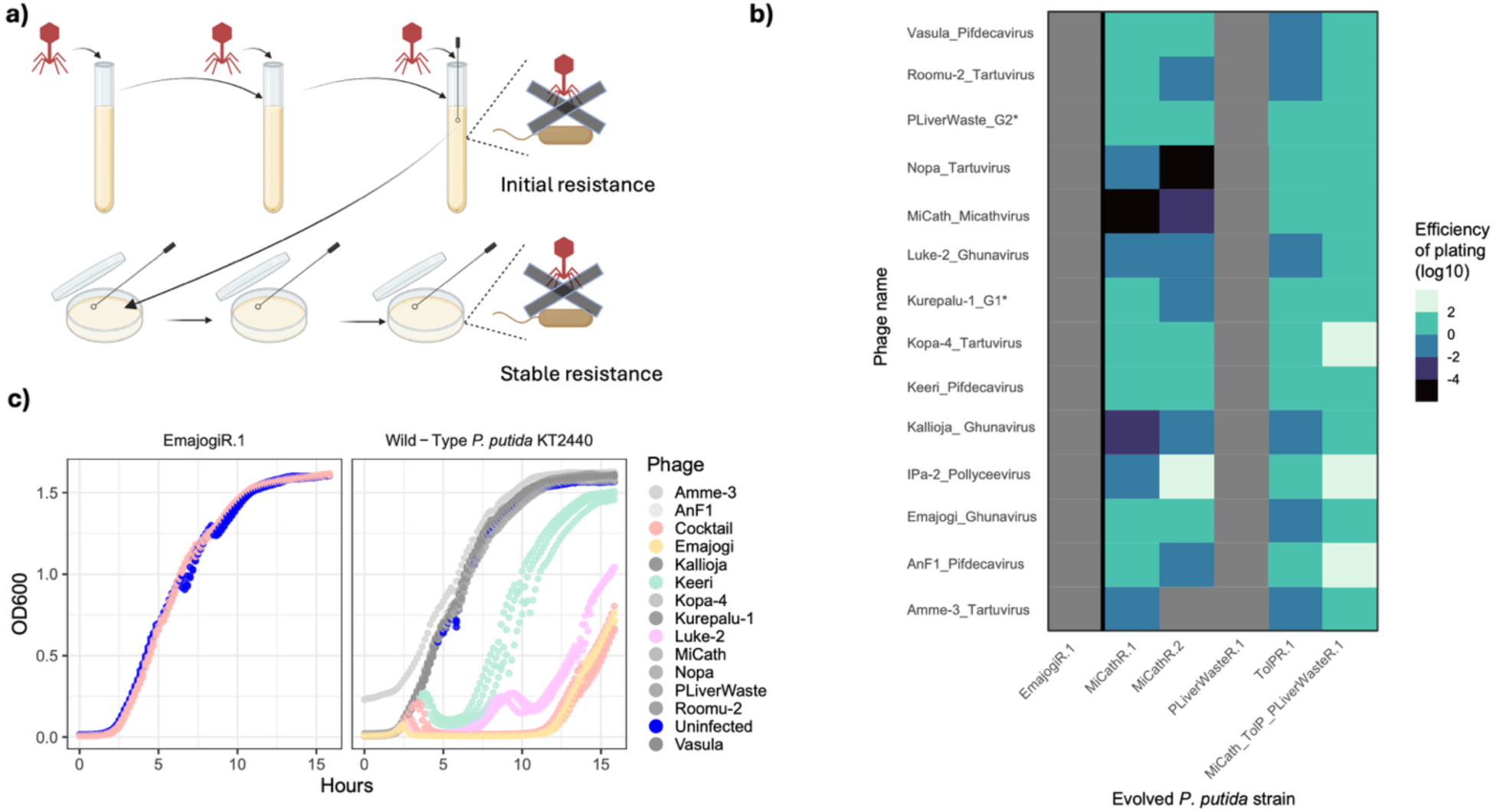
Experimental evolution yields strains stably resistant to multiple genera of *P. putida* phages. **a)** Schematic of experimental evolution. “Initial resistance” was tested after overnight incubations with phage additions. If initial resistance was found, the strains were passaged three times without the presence of phage, and phage resistance was tested again to determine “stable resistance”. **b)** EOP for a diverse phage panel compared to the parent strain noted below. Grey boxes indicate to no plaques detected at all concentrations tested. G#* is currently not a defined genus at ICTV^39^. **c)** Liquid cultures were infected at an MOI of 0.2 and OD_600_ was measured post infection. *P. putida* KT2440 was tested with each individual phage and a sixteen- phage cocktail, phages are colored coded. EmajogiR.1 was tested only with the sixteen-phage cocktail. The sixteen-phage cocktail was comprised of an equal concentration of each phage (0.02:1 virus to cell).

### Genetic basis of EmajogiR.1 phage resistance

To identify the genetic mechanism of phage resistance for Emajogi.R1, we analyzed DNA and RNA sequencing. Comparing assembled genomes of EmajogiR.1 and our in-house wild-type KT2440 revealed that they differed by a single frameshift deletion, which reverted the early stop codon in the *aprD* gene of a T1SS operon (**Figure 3a and 3b**). The T1SS operon contains three genes: *aprD*, an ATP binding cassette transporter protein, *HylD*, the membrane fusion protein, and an outer membrane factor (*OMF*)^18^. The same mutation was observed in PLiverwasteR.1, which evolved independently from *P. putida* mt-2 but displays the same broad phage resistance (**Figure 3c, Figure 2b**).

**Figure 3.**
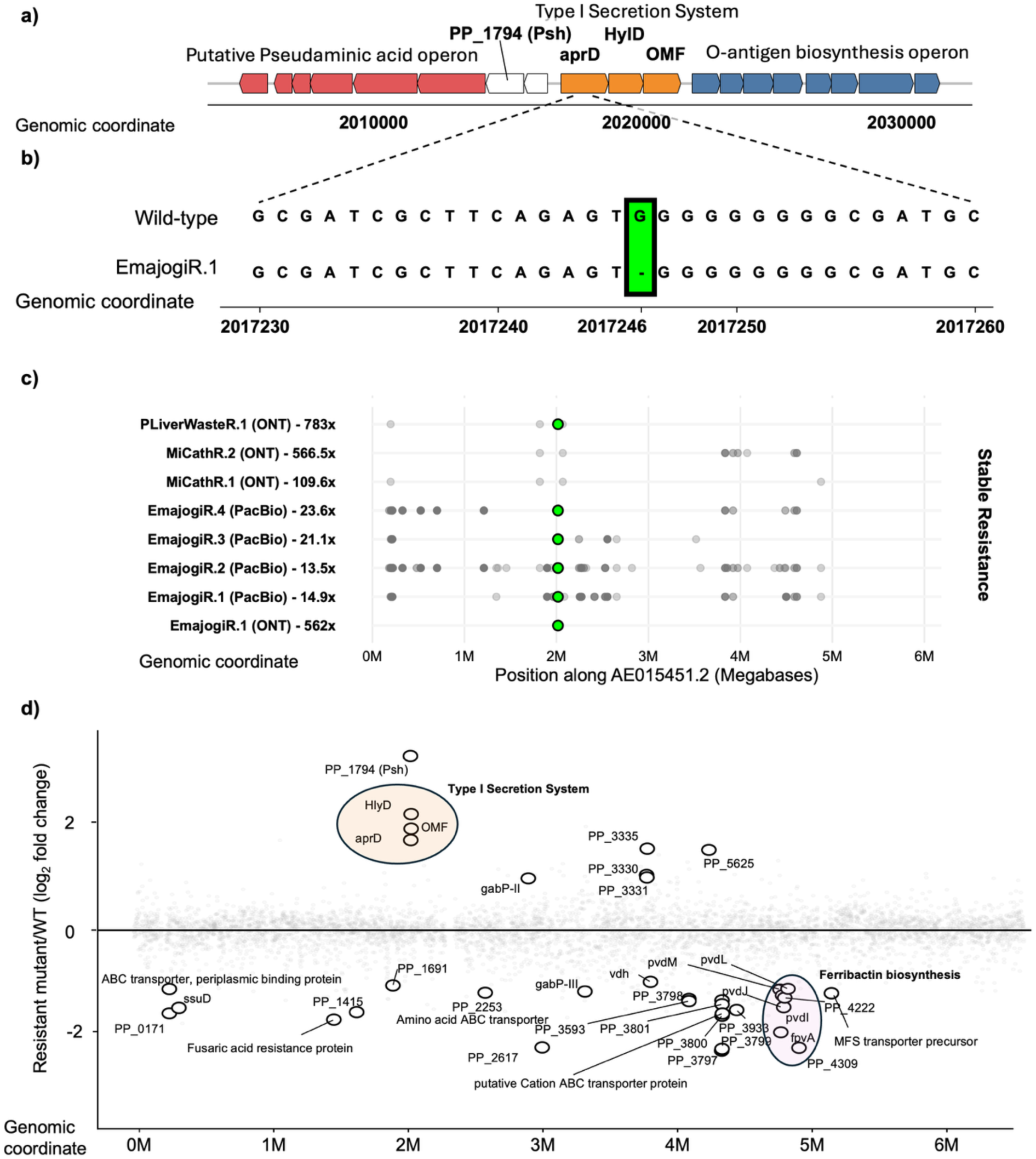
EmajogiR.1 has a single point mutation that restores the aprD ATPase in a T1SS and several transcriptionally upregulated operons. **a)** Genetic region of T1SS and *Psh*. Any gene found significantly upregulated is in bold. **b)** Nucleic acid sequence of *aprD* gene from the T1SS. The top sequence is wild-type *P. putida* KT2440, which has an early termination codon, classifying it as a pseudogene. The bottom shows EmajogiR.1 sequence with a frameshift deletion restoring functionality of the aprD ATPase subunit of the T1SS operon. **c)** The mutation analysis of each evolved resistant strain is shown (rows). The name of the strain, sequencing technology (ONT is oxford nanopore), and genome coverage are listed. Each dot represents a polymorphism, deletion, or insertion in the final assembled genome after subtracting out mutations found in the in-house parental strain the mutant was propagated from. EmajogiR strains were propagated from *P. putida* KT2440, while all other strains were propagated from *P. putida* mt-2. Green dots mark single-nucleotide deletions found between nucleotides 2,017,246 to 2,017,254, which lies within the *aprD* gene of the incomplete T1SS in *P. putida* KT2440. Note that all stably resistant strains that were completely resistant against the phage panel have a single nucleotide deletion in the *aprD* region. When sequencing coverage is low, more mutations were observed, which may represent assembly artifacts. **d)** Transcriptional fold change across the genome of EmajogiR.1 compared to *P. putida* KT2440. The T1SS operon, which resides next to Psh and contains the SNP in aprD is highlighted in orange. The downregulated longest operon, coding for ferribactin biosynthesis, is highlighted in pink.

Since only one nucleotide differed in our DNA sequences, we evaluated transcriptional changes as well to understand the mechanism of phage resistance. Differential expression analysis revealed several operons altered in EmajogiR.1 relative to wildtype KT2440 (**Figure 3d**). The longest differentially expressed block contained ferribactin/pyoverdine related genes (pvdI, pvdJ, pvdL, pvdM, fpvA). The most strongly upregulated transcript (9.9fold) encoded a hypothetical protein, Psh. The *Psh* gene is located within a putative pseudaminic acid/O-antigen capping operon and is directly opposite the restored T1SS which was also upregulated (**Figure 3a and 3d**). Because T1SSs typically sit adjacent and antisense to the substrate they export^19^, we hypothesized that this restored T1SS transports Psh to the outer membrane, where it has a role in phage defense.

### EmajogiR.1 phage resistance requires a restored T1SS and Psh

We tested whether the restored T1SS and the adjacent Psh are required for phage defense by overexpressing them in wildtype *P. putida* KT2440, which normally lacks this restored T1SS, and challenging with Emajogi. Overexpressing Psh alone provided mild phage resistance (**Figure 4a**). In contrast, only overexpressing introducing the full T1SS (aprD, HylD, and OMF) from EmajogiR.1 conferred strong protection (**Figure 4b**), whereas overexpressing the native pseudogene version of the system did not (**Figure 4c**). Overexpressing both Psh and the restored T1SS in conjunction conferred complete resistance comparable to EmajogiR.1 (**Figure 4d**). Overexpressing the pair also conferred MiCath resistance in *P. putida* S12, despite *P. putida* S12 lacking a T1SS and Psh homolog (**Figure 4e**).

**Figure 4.**
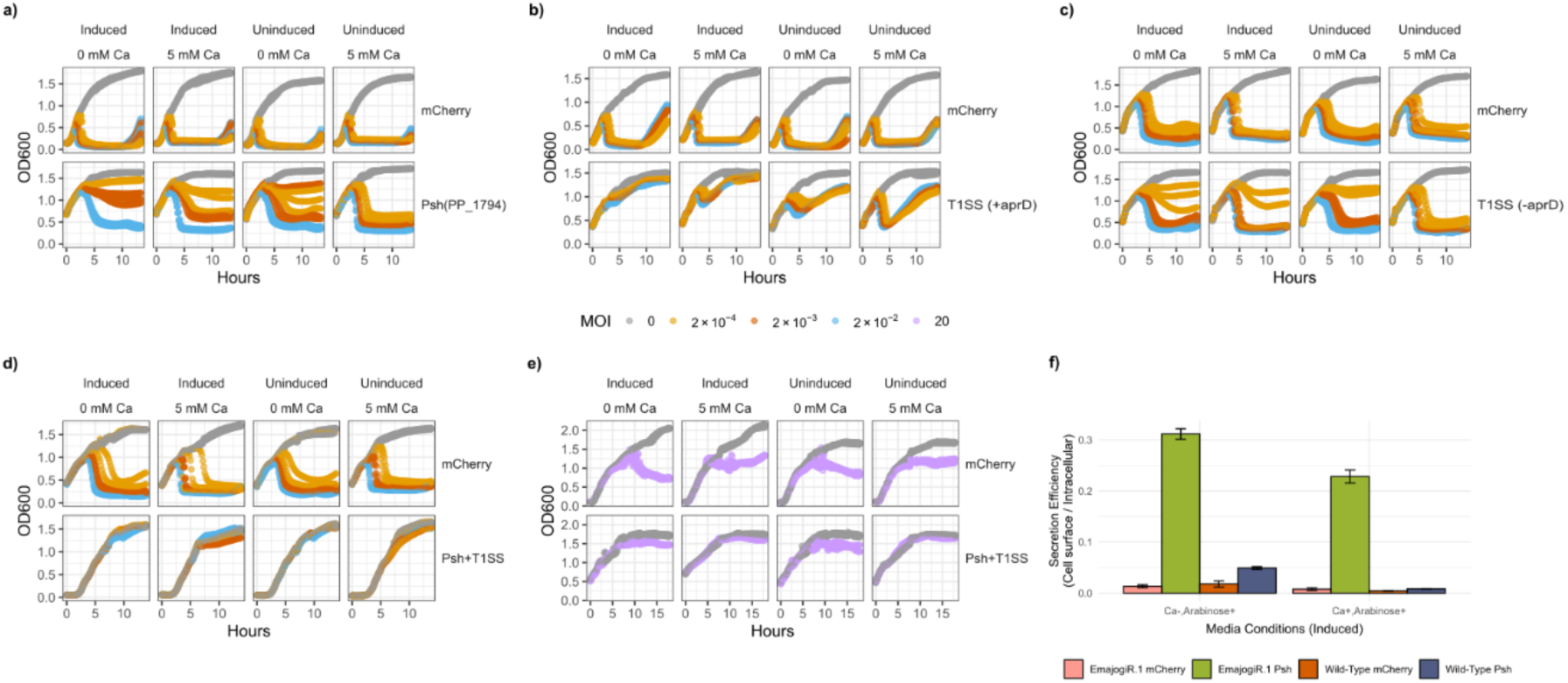
Psh protein is secreted by restored T1SS and provides phage resistance. Growth curves of *P. putida* KT2440 infected with different MOIs of Emajogi, containing plasmids with arabinose-inducible promoters, overexpressing **a)** Psh protein, **b)** restored T1SS (found in EmajogiR.1, with a full-length aprD subunit), **c)** defective T1SS (found in KT2440, with an early termination codon in the aprD subunit), and **d)** Psh protein + restored T1SS. MOIs are differentially colored and the key is shown in the center of the figure. **e)** Growth curve of *P. putida* S12 strain infected with phage MiCath at MOI=20, overexpressing the complete Psh/T1SS system. For each strain, induced and uninduced conditions are shown with and without the addition of 5mM CaCl_2_. **f)** Export efficiency of induced HiBit-tagged Psh protein and mCherry expressed in KT2440 and EmajogiR.1 with and without CaCl_2_.

We sought to determine if the restored T1SS exports Psh to the outer membrane to prevent phage infection. A N-terminal HiBit tagged Psh showed more than 6-fold higher export in EmajogiR.1 compared to wild-type KT2440 (**Figure 4f, Supplemental Figure 1**). We then sought to understand if the exported Psh could be protective to other cells in a co-culture. We used the EmajogiR.1 strain and a mCherry labeled KT2440 with no cross-protection evident (**Supplemental Figure 2**). This indicates the defense is not shared between cells in close contact with each other. Lastly, we examined LPS fraction of *P. putida* KT2440 and EmajogiR.1 to understand if there were any changes to the LPS. No differences in LPS were observed between wild-type *P. putida* KT2440 and EmajogiR.1, but the total protein profiles of the same extraction showed a large aggregate only present in EmajogiR.1 (**Supplemental Figure 3**). Altogether, these experiments suggest that the restored T1SS exports Psh.

### EmajogiR.1 has no fitness defects except reduced pyoverdine production

To test if EmajogiR.1 could still be beneficial for traditional biomanufacturing processes, we tested its ability to grown on a variety of hydrolysates, which represented typical and state of the art economical carbon sources for growing high biomass of *P. putida,* to directly compare biomass conversion capabilities of wild-type *P. putida* KT2440 and EmajogiR.1. We observed that EmajogiR.1 retained its phage resistance and was able to utilize the carbon sources as efficiently as wild-type *P. putida* KT2440 (**Figure 5**), confirming that this mode of phage resistance was insensitive to carbon type.

**Figure 5.**
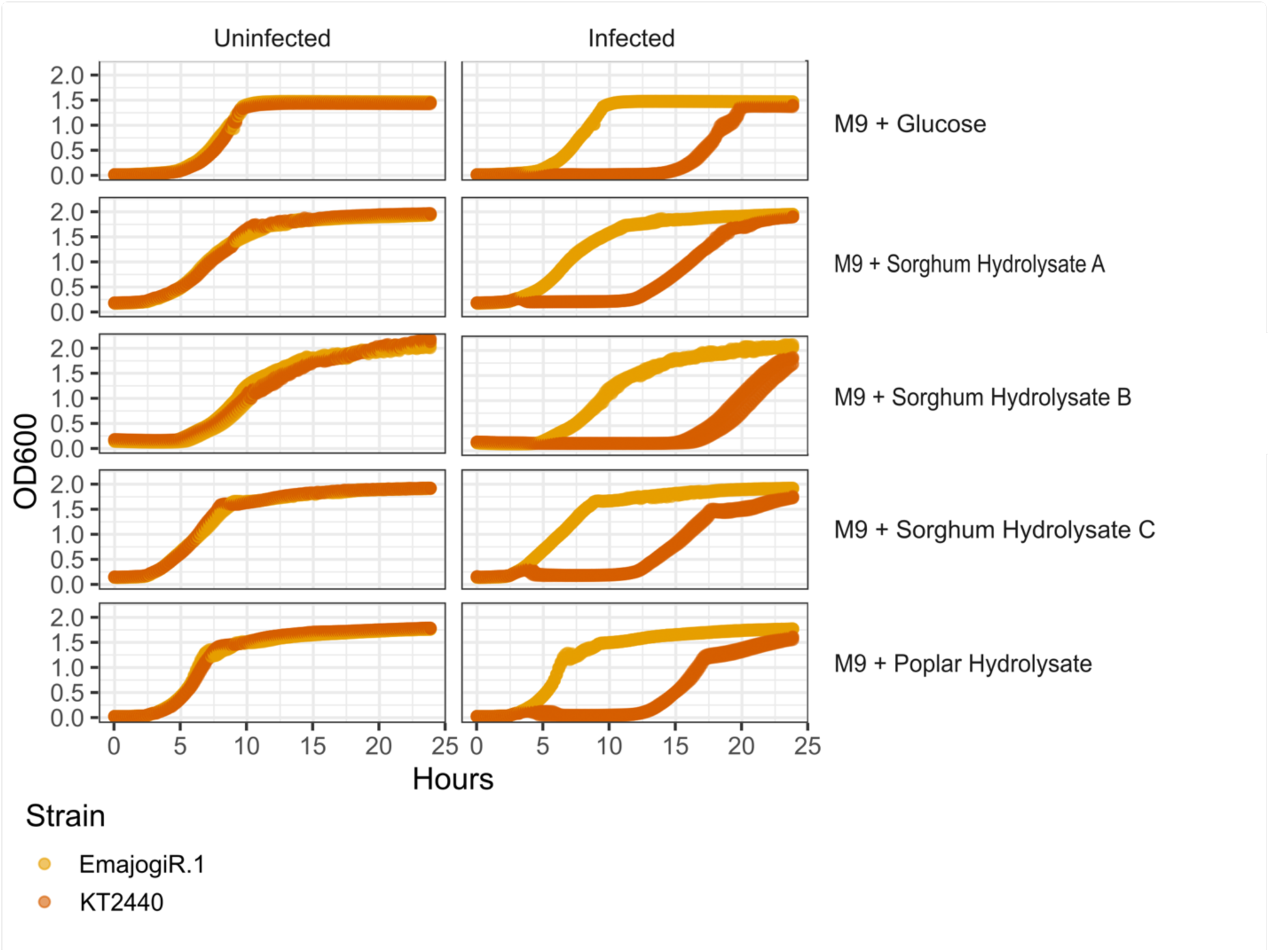
No growth difference is observed between *Pseudomonas putida* EmajogiR.1 and *P. putida* KT2440 when grown in a variety of carbon sources. *P. putida* KT2440 or EmajogiR.1 were grown in M9 media supplemented with the indicated carbon sources. Uninfected cultures are in the left column, and Emajogi infected culture (MOI = 0.002) are in the right column.

We observed no difference in electroporation efficiency of EmajogiR.1 and wild-type *P. putida* KT2440 (**Supplemental Figure 4**). We additionally observed no minimum inhibitory concentration (MIC) differences for a suite of popular antibiotics (Gentamicin, Kanamycin, Chloramphenicol, Tetracycline, Carbenicillin, and Trimethoprim) (**Supplemental Table 1**). Taken together, this suggests EmajogiR.1 is capable of transformation and has no antibiotic bias that would reduce its effectiveness in biomanufacturing applications.

Thus far, no observable phenotype tradeoff was evident, which was atypical for a system conferring phage defense to numerous phages^20^. The RNA sequencing revealed downregulation of multiple pyoverdine biosynthesis genes (**Figure 3d**). We observed a significant reduction in pyoverdine biosynthesis with no change in bacterial growth (**Supplemental Figure 5**). Pyoverdine is an iron-chelating small molecule which has been shown to increase seed colonization by *P. putida*^21^, as well as increase virulence in pathogenic *Pseudomonas* strains^22^. However, reduction of pyoverdine in biomanufacturing *Pseudomonas* species is considered beneficial since it can improve metabolism and fluorescent measurement sensitivity^23^.

### Diverse bacterial systems carry the Psh defense system

We sought to assess the prevalence of this novel phage defense system across bacteria. Psh is annotated to have two domains: a helix rich DUF4214 domain at the C-terminus and an N-terminal RTX-like repeat domain, which is supported by the predicted 3D structure (**Figure 6a**). Psh was queried against the UniRef90 database revealing hundreds of homologs spanning either the full-length protein or only the C-terminal DUF4214 domain (**Figure 6b**). Outside of *Gammaproteobacteria,* nearly all homologs corresponded solely to DUF4214-containing proteins, whereas *Gammaproteobacteria* encoded for both DUF4214-only proteins and RTX– DUF fusions resembling Psh from wild-type *P. putida* KT2440 (**Figures 6b and 6c**). The RTX–DUF fusion architecture present in *P. putida* KT2440 therefore appears conserved within closely related *Gammaproteobacteria* (**Figure 6c**). To identify what other domains Psh-related proteins contained, we queried the predicted functional domains of genes encoding the DUF4214 domain which did not contain an RTX-like domain, and observed diverse domains, including secreted peroxidases, peptidases, and immunoglobin-like folds (**Supplemental Figure 6**).

**Figure 6.**
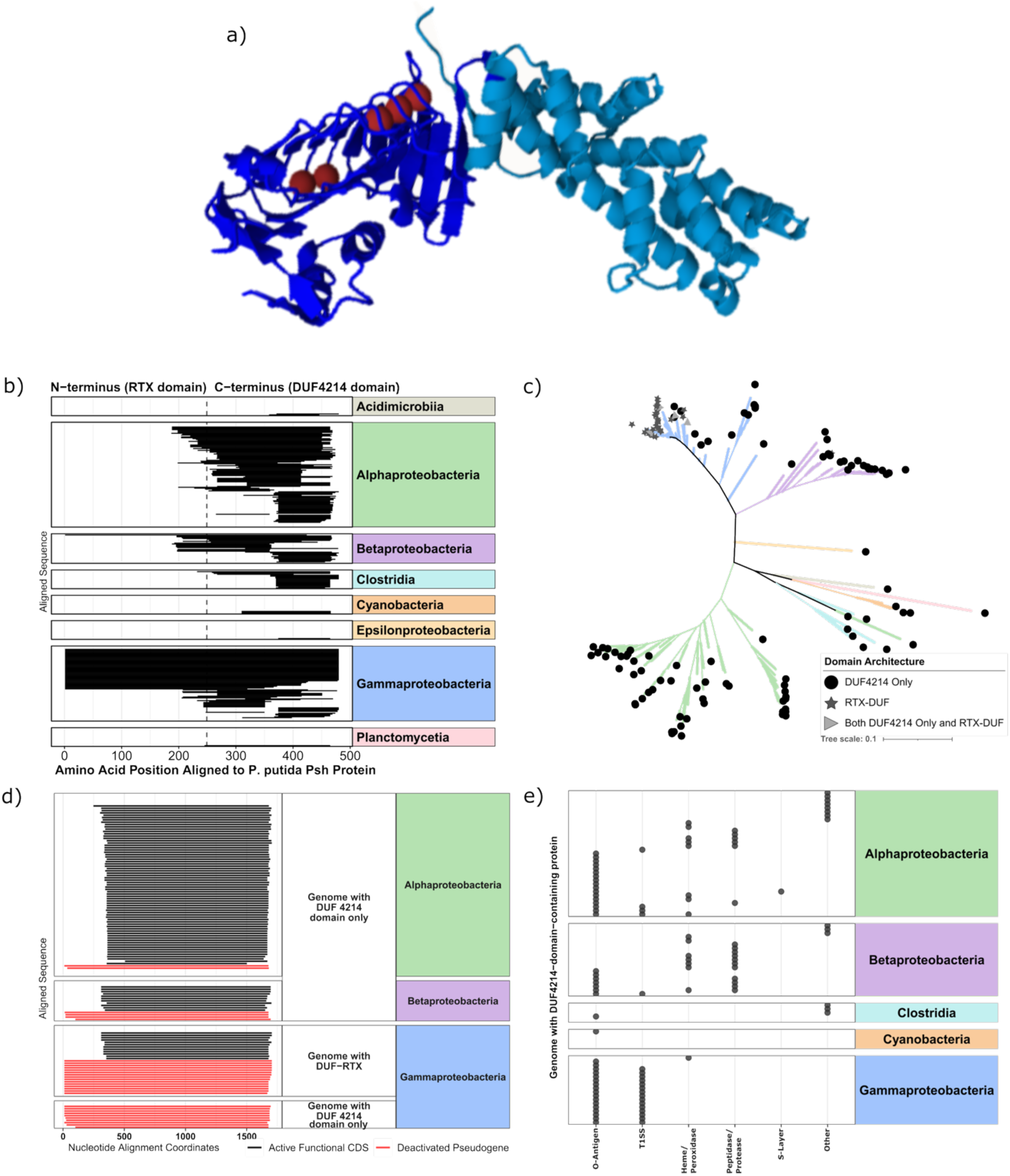
The Psh protein and associated T1SS are widespread among diverse bacteria. **a)** Representative three-dimensional structural model of the Psh protein. N-terminal RTX-repeat domain is dark blue and the C-terminal helix-rich DUF4214 domain is teal. Solid red spheres map to predicted calcium-binding pockets. **b)** Local sequence alignments mapping Psh to homologous proteins found in diverse bacteria. The taxonomy of each bacterial class is shown and color-matched to the corresponding tree shading in (**c**). Dashed vertical line signifies approximate C-terminus and N-terminus domain spits. **c)** Unrooted maximum-likelihood phylogenetic tree of representative bacterial 16S rRNA gene sequences that carry the DUF4214 locus (n = 244 unique strains). Overlaid terminal geometric icons indicate if the host genome contains a gene with DUF4214 only, the full length Psh (DUF4214 and RTX-repeat domain) or both proteins. **d)** Each row represents a T1SS *aprD* gene found in a unique genome through translated sequence searches. Protein where the KT2440 pseudogene (+1 frame coding for *aprD* pseudogene) is found are highlighted in red. Black are proteins where only full-length *aprD* gene (+2 frame) are found. The bacterial class and what type of Psh protein is shown on the y-axis. **e)** Gene neighborhood co-occurrences within a rolling 10 kbp window flanking the *DUF4214* gene. Each horizontal row represents a unique genomic locus instance. Solid charcoal markers map presence of six core companion frameworks.

Since Psh requires a T1SS to be secreted (**Figure 4**), we next examined the distribution of the pseudogenized versus full-length T1SS *aprD*. Using the naturally occurring frameshifted pseudogene (*aprDB)* as a query, we found that the truncated T1SS is common among *Gammaproteobacteria* and occurs across a broader bacterial phylogeny than Psh (**Figure 6d**). Having established that RTX–DUF fusion proteins and associated T1SS are broadly distributed across bacteria, we next investigated neighboring genes to gain additional insight into their potential functions.

In *P. putida* KT2440, the *Psh* gene sits between the O-antigen capping and biosynthesis operon and the T1SS (**Figure 2a**). This genomic context is largely conserved across *Gammaproteobacteria* (**Figure 6e**). However, in *Alphaproteobacteria* and *Betaproteobacteria* the surrounding operons are more diverse (**Figure 6e**). Yet, across all bacterial classes surveyed, over 80% of Psh proteins detected occur adjacent to O-antigen biosynthesis operons or T1SS. Therefore, we sought to understand if there were any predicted interactions between Psh and O-antigens. Structural modeling predicted that Psh can bind the O-antigen and its putative pseudaminic acid cap at nanomolar efficiency (**Supplemental Figure 7**). Taken together, this suggests that there is a conserved genetic organization which may predict the relevant binding partners for Psh.

## Discussion

Here we characterized a novel, broad-spectrum phage-resistance mechanism in the biomanufacturing strain *Pseudomonas putida* KT2440 (**Figure 1**). This system comprises a previously uncharacterized RTX-DUF4214 fusion protein, designated Psh, which is exported by a restored T1SS, and binds to the O-antigen of the LPS. This novel mechanism conferred resistance to sixteen different phages, spanning six distinct genera likely by masking their receptor. There was no growth defect observed across a variety of carbon sources, no defects in transformation efficiency or protein production. The only fitness cost observed was a reduction in pyoverdine production. Further, Psh homologs were conserved across *Gammaproteobacteria* and consistently found near T1SS and O-antigen operons.

Discovery of phage-resistance mechanisms has exploded in recent years, partly due to the resurgence of phage therapy^24^. In phage therapy applications, there is a need to understand and overcome the bacterial defense; however, for many non-therapeutic applications, these defense mechanisms may be beneficial. Biomanufacturing is rapidly growing due to advances in synthetic biology making difficult-to-source products readily obtainable from engineered microbes. While these processes are highly tuned, they occasionally suffer from phage contamination, leading to extended downtime, costly cleanup efforts, and loss of valuable product^4^. Leveraging the known phage-resistance mechanism for biomanufacturing bacteria could remove this problematic hurdle.

One of the main reasons that phage defense mechanisms have not yet been widely employed to bolster biomanufacturing strains is the unknown and unwanted fitness costs associated with expression of these systems^20^. In the system described here, we show no fitness impacts except for reduced pyoverdine production. Reduced pyoverdine production could impact survival in low iron rich environments, such as soil, where *P. putida* is naturally found. While in soil and root environments reduced pyoverdine does impact niche-establishment^21^, swarming, and root colonization, this is not the case in biomanufacturing^22^. Pyoverdine is seen as a contaminant and inhibitor in biomanufacturing since it can bind iron with high affinity, induce biofilm formation, and interfere with fluorescent protein analysis. Overall, the reduction in pyoverdine in this strain should make the strain more attractive for biomanufacturing.

The current hypothesized model is that the Psh protein masks the receptor these phages use to inject their DNA into the cell (**Figure 1**). We show evidence that Psh is exported by the cell only when the *aprD* gene is full length resulting in a functional T1SS and the two- component (functional T1SS and Psh) system can confer resistance in strains that do not contain this system. While we know this system is working at the cell surface, we cannot determine exactly how Psh interacts with the O-antigen. This may be as aggregates that surround the O- antigen or through a tight binding association that is further enhanced by calcium ions. Further, we show there is resistance conferred to phages whose receptor is not the LPS (AnF1, Vasula, and Emajogi)^16^ suggesting this mechanism is a receptor masking mechanism. While a few other secreted protein receptor masking mechanisms are known (TraT in *Escherichia coli*^25^, RsaA in *Caulobacter vibioides*^26^, and cell-binding protein A in *Staphylococcus aureus*^27^) these mechanisms do not bind to the O-antigen.

There is a broad phylogenetic span of proteins similar to Psh, however most of these proteins, outside of *Gammaproteobacteria*, lack the RTX-domain. The RTX domain is located adjacent to the C-terminal secretion signal and is where calcium residues bind to stabilize the secreted protein. This suggests that this type of mechanism may exist in other bacteria but relies on another protein domain for these functions. While the DUF4214 domain is found in nearly all classes of bacteria examined, the function is poorly characterized. Based on the data shown here, this domain likely has a role in stabilization of the complex on the pseudaminic cap. The gene neighborhood of Psh is typically located near the O-antigen and T1SS in *Gammaproteobacteria* but not in other bacteria classes examined suggesting the transport mechanism may only exist in this class. Overall, this mechanism is likely useful for protection from phage in other Gram-negative bacteria. We have uncovered a novel strategy *P. putida* uses to evade phage infection. This strain has no defects in traditional biomanufacturing metrics (growth, transformation, protein production) and has reduced pyoverdine production which benefits biomanufacturing processes.

## Methods

### Phage propagation

Phages were propagated using double agar overlays^28^. A single plaque was picked and resuspended in 100 μL SM buffer (50 mM Tris-HCl (pH 7.5), 100 mM NaCl, and 8 mM MgSO₄ ⋅ 7H₂O), mixed with 125 μL of overnight culture of *P. putida* and 4 mL of LB top agar (0.5% agar), and plated onto a fresh LB agar plate. Web plates were then flooded with 8 mL of SM buffer, scraped, spun at 3000xg for 5 minutes to remove excess agar, then filtered through a 0.2 μm filter. Titers of phage lysates were quantified via spot assay.

### *P. putida* phage resistance evolution

*P. putida* mt-2 cultures were grown overnight in 25 mL of LB broth at 30°C with shaking at 230 rpm. Overnight cultures were diluted to an OD_600_ of 0.05 and grown to an OD_600_ of 0.2 when individual bacteriophages, MiCath, PLiverWaste, Tolp, and the phage cocktail consisting of equal proportions of all three phages, at MOIs of 0.1, 1, and 5 were added.

Samples were collected at 3-hour intervals for a total of three sampling time points each day. Immediately following sample collection, bacterial growth was quantified by measuring the OD_600_ and viable bacterial counts were determined by serial dilution spot plating at 30°C. Following the final sampling time point, the cultures were incubated overnight, and process was repeated daily up to seven consecutive days to facilitate host–phage co-evolution. Final cultures were streaked three times on LB agar to remove any phage particles.

To produce the Emajogi resistant strains, Emajogi was infected in a 96 well plate at a 0.001 MOI, any culture that had bacterial growth after 24 hours was streaked three times on an LB agar.

### Efficiency of Plating Assay

*P. putida* KT2440, mt-2, and all phage-resistant strains derived from the experimental evolution were grown overnight in LB media. Sixteen phages from six distinct genera were serially diluted in phage buffer to 10^-11^. Each overnight *P. putida* strain was diluted 1:6 in LB top agar, then plated onto the LB plate. After the top agar layer was cooled and dried, 2uL of each serially diluted phages were plated onto the top agar and incubated at 20°C. EOP was calculated against their respective parent strains using the following equation:

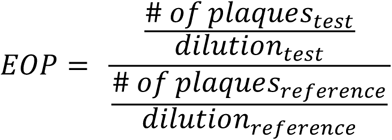

### DNA extraction and sequencing preparation

Sample sequenced using Pacific Biosciences (PacBio) platforms were prepared as follows. DNA was extracted from cells using the Nanobind kit protocol (Pacific Bioscience, cat # 102-207-600; cat# 102-207-700) according to the manufacturer’s instructions. Cell pellets were lysed with Proteinase K and lysis buffers, followed by RNase A treatment. DNA was captured on a Nanobind disk in the presence of isopropanol, washed with wash buffers, and eluted in Buffer LTE. The eluate was incubated overnight at room temperature to ensure complete DNA solubilization before quality assessment and downstream applications.

DNA was sheared using a g-TUBE (Covaris, cat. # 520079)) according to the manufacturer’s instructions. A 150 µL of fully dissolved particle-free DNA was loaded into the g-TUBE and centrifuged at 4800xg to transfer the sample through the shearing tube. DNA libraries were prepared using the SMRTbell Express Template Prep Kit 3.2 (Pacific Biosciences) following the manufacturer’s instructions. Final libraries were quantified using a Qubit Flex fluorometer and assessed for fragment size using an Agilent 4200 TapeStation. The samples were sequenced on a Sequel IIe System (Pacific Biosciences) according to the manufacturer’s protocol.

Sample sequenced using Oxford Nanopore Technologies (ONT) were sent to plasmidsaurus where DNA and sequencing libraries were prepared.

### De Novo Genome Assembly and Coverage Evaluation

Long-read sequencing data generated via ONT and PacBio platforms were assembled *de novo* using the Flye assembly pipeline^29^ (version 2.9). For each isolate, raw reads were processed utilizing default long-read optimization parameters to generate unfragmented contig files. Quality metrics, structural configurations (circularity), and individual contig coverage values were extracted directly from the compiled assembly_info.txt data outputs generated within each strain-specific directory. Whole genome sequencing confirmed the cultures were uncontaminated, with over 95% of DNA reads aligning to *P. putida* KT2440.

### Variant Identification and Comparative Genomics

Assemblies were evaluated using the MUMmer comparative genomics package^30^ (version 3.23) to identify single-nucleotide polymorphisms (SNPs) and short insertions/deletions (indels) across the evolution experiment strain. Individual assembled mutant genomes were mapped against the universal *Pseudomonas putida* KT2440 public reference standard genome (AE015451.2) using the nucmer alignment engine. To bypass large structural assembly discrepancies and false-positive calls arising from repetitive regions, alignments were executed utilizing the global mapping mode (--maxmatch). To isolate experimental mutations from pre- existing background mutations, long-read in-house parent sequences were subtracted from their respective descendant mutant profiles.

### Subpopulation Validation

To investigate whether the single-nucleotide deletion observed within the *Pseudomonas putida* KT2440 homopolymer region represented a biological subpopulation or long-read sequencing artifact, a short-read framework was used. Paired-end Illumina sequencing tracks from the control wild-type *P. putida* KT2440 sample EmajogiR.1 were compared.

Raw paired-end short reads were mapped to the reference *P. putida* KT2440 complete genome sequence (AE015451.2) using the short-read preset mode (-ax sr) of minimap2^31^ (v2.30). To ensure maximum statistical power over the homopolymer target locus, alignments were processed at full sequencing depth without read downsampling. The resulting raw sequence alignment map tracks were progressively converted to BAM format, coordinates- sorted, and indexed using samtools^32^ (v1.23). A 20-base pair window spanning genomic coordinates 2,017,241 to 2,017,261 was selected for direct locus inspection. Quantitative variant evaluation was conducted using samtools mpileup coupled with the reference sequence template. A mapping and base quality threshold of -Q 25 was applied to the pileup stream to completely exclude ambiguous or low-confidence alignment records.

The subpopulation fraction (%) for each sample was resolved as the ratio of deletion events relative to the total number of high-confidence evaluated reads overlapping the coordinate.

### Transcriptomics

Triplicate cultures of *P. putida* KT2440 and the evolved strain EmajogiR.1 were subcultured into LB and grown for 3 h at 30 °C in 14 mL culture tubes without amendments. Cultures were preserved directly at a 1:2 (culture:preservative) ratio using DNA/RNA Shield (Zymo Research, cat. # R1100-50) without centrifugation and stored at −80 °C until processing. Samples were treated with the RNase-Free DNase I Kit (Norgen Biotek, cat. # 25710) prior to RNA extraction.

Total RNA was isolated using the Quick-RNA Fungal/Bacterial Miniprep Kit (Zymo Research, cat #. R2014) following the manufacturer’s protocol. RNA libraries were prepared and sequenced by SeqCoast using Illumina TruSeq Stranded RNA-seq chemistry.

Reads were mapped to the *P. putida* KT2440 reference genome using Bowtie^33^ (v2.3.0). Gene-level counts were generated using featureCounts, and differential expression was performed with DeSeq2^34^. All downstream processing and visualization were performed in RStudio.

### Phage Defense in Liquid Culture

Plasmids for Psh, the T1SS containing the aprD pseudogene and restored gene, and mCherry were cloned into the pJLCKan backbone^35^ using Gibson assembly (Primers in **Supplemental Table 2;** Plasmids listed in **Supplemental Table 3**). These plasmids were transformed into *P. putida* KT2440 or P. putida S12 and grown overnight in LB supplemented with kanamycin (50 µg/mL) shaking at 30°C. The overnight cultures were subcultured 1:50 into LB supplemented with kanamycin (50 µg/mL) with one of the following inducers 0% arabinose/0mM CaCl_2_, 0.2% arabinose/0mM CaCl_2_, 0% arabinose/5mM CaCl_2_, and 0.2% arabinose/5mM CaCl_2_ and grown for 4-5 hours shaking at 30°C. 200uL of KT2440 strains from each growth condition was infected with 2uL of Emajogi at an MOI of 2×10^-2^ to 2×10^-4^. 200uL of S12 strains from each growth condition was infected with 2uL of MiCath at a MOI of 20. Growth curves were assayed overnight using OD_600_ measurements in a Tecan Spark plate reader with OD_600_ readings measured every 10 minutes while shaking at 30°C. To confirm that cultures were expressing their constructs, mCherry (excitation 585nm/emissions 615nm) fluorescence was measured.

### Plasmid and transformation conditions

Plasmids were transformed into *E. coli* DH5alpha and then into *P. putida* strains. *P. putida* electrocompetent cells were prepared by subculturing an overnight incubation grown in LB media 1:50 then growing for an additional 3-5 hours. Cells were chilled on ice and washed by spinning down 5 minutes at 5000xg at 4°C, resuspending in pre-chilled 10% glycerol 3 times, and finally resuspending in 1/50 original volume 10% glycerol, aliquoted into 50 ul portions, and frozen at -80C. To transform, cells were thawed on ice for 5-30 minutes, then ∼40-150 ng of plasmid DNA was added. This mixture was transferred to pre-chilled 1 mM electroporation cuvettes (Thermo Fisher P41050) and shocked with the capacitance at 25 µF, the resistance at 200 Ω, and the voltage at 1.8 kV. Cells were recovered with LB and plated on LB agar supplemented with kanamycin (50 µg/mL) after 1-2 hours recovery.

### Psh localization assay

A HiBit tag was added to the N-terminus the Psh protein on an overexpression plasmid. This vector was transformed into wild-type *P. putida* KT2440 and EmajogiR.1 strains. Overnight cultures of each strain were picked in LB without inducers or CaCl_2_. The next day, cultures were diluted into LB with/without calcium and with/without 0.2 % arabinose and grown for 3 hours. For the intracellular and cell-surface fractions, 10 μL was aliquoted into a 384 well black plate with a black flat bottom. For the supernatant fraction, 150 μL of each condition was filtered (5 minutes, 1000xg) in a MultiScreen GV filter plate (Millipore Sigma, cat. # MAGVS2210), and 10 μL was also aliquoted into the same 384 well black plate.

Next, working buffers of Nano-Glo® HiBiT Extracellular Detection System (Promega, cat. # N2420) and Nano-Glo® Luciferase Assay System (Promega, cat. # N1110) were prepared using ½ of the recommended concentration of LgBiT protein enzyme unit as recommended in each. The extracellular system was used to detect the supernatant and extracellular fractions, while the Luciferase system was used to detect the intracellular fraction. Once these buffers were prepared, 10 μL was added to the black well plate and immediately placed in a plate reader. To capture the luminescence profile of the NanoLuc reporter, a spectral scan was performed across an emission wavelength range from 400 nm to 560 nm. Signal collection at each wavelength step was acquired with an integration time of 500 ms and a 0 ms settle time. Raw luminescent data points across the scanned spectrum were collected and expressed as counts per second. NanoLuc® luminescence was monitored in kinetic mode over a total duration of 40 minutes, with luminescent measurements acquired at 2-minute intervals. To maintain sample homogeneity throughout the kinetic run, the microplate was subjected to double-orbital shaking for 5 seconds prior to each measurement point at a frequency of 150 rpm and an amplitude of 2 mm.

The OD_600_ of each condition was taken on a sperate plate reader shortly after starting the luminescence scan and used for biomass normalization.

### SDS-PAGE of LPS and Total Protein

Wild-type *P. putida* KT2440 and EmajogiR.1 strains were grown in LB media at 30°C shaking at 250 rpm overnight. Two 100uL aliquots from each strain were harvested via centrifugation at 1000xg for 5 minutes. Supernatants were discarded, and the pellets were washed twice with 500uL of PBS pH 7.2. Samples were spun down again at 1000xg for 5 minutes. One supernatant aliquot from each strain was collected for analysis via SDS-PAGE. The other aliquot was treated with 500uL of 5mM EDTA for 30 minutes at room temperature. The EDTA treated samples were spun down at 1000xg for 5 minutes, then the supernatants were collected for analysis via SDS-PAGE.

SDS-PAGE was performed at 120V/400mAmps for 30 minutes on 12% Mini-PROTEAN TGX precast gel (Bio-Rad, cat. #4561044). Resulting gel was processed for LPS and total protein staining using Pro-Q Emerald 300 LPS stain kit (ThermoFisher, cat. # P20495) and SyproRuby stain kit (ThermoFisher, cat. # S12000) respectively following the manufacturer’s protocols. The stained gel was imaged using the AzureBiosystems imager.

### Media preparation of hydrolysates

The biomass hydrolysates were gifted by the DOE Joint BioEnergy Institute and conditioned for microbial cultivation. The hydrolysates were diluted 1/4 into 1x M9 without carbon and supplemented with iron (30.00 µM ferric ammonium citrate, 31.20 µM citric acid) and trace metals ((46.26 µM H₃BO₃, 9.15 µM MnCl₂, 1.61 µM Na₂MoO₄, 0.77 µM ZnSO₄, 0.32 µM CuSO₄, 0.17 µM Co(NO₃)₂). As a positive control 0.4% glucose final concentration was added to the 1x M9 described above in place of the hydrolysate.

### Pyoverdine production

To assess if a culture produced pyoverdine, overnight uninduced cultures without calcium of both wild-type *P. putida* KT2440 and EmajogiR.1 with pJLCKan/mCherry were subculture in uninduced LB media without calcium, grown for 4 hours, and aliquoted into 200 μL portions and placed inside a plate reader. Pyoverdine abundance was measured by exciting 400 nm (± 10 nm) and measuring emission at 460nm (± 15 nm).

### Sequence Homology Analysis

To find homologs of the Psh protein, the Uniref90 dataset^36^ was downloaded and queried in a local database with Diamond (v2.2.4)^37^ using the blastp command with the –sensitive option. Matches with bitscores below 50 were removed, and the top two hits at the species level were kept reducing graphing closely related strains. To query the taxonomic lineage of the bacteria harboring these Psh homologs, a 16S alignment of NCBI taxonomic IDs of bacteria with Psh proteins was assembled with IQTree (v3.1.2)^38^ using default settings. To find homologs of the truncated *aprD* gene in other bacteria, the *aprD* pseudogene from wild-type KT2440 was used to search RefSeq bacteria (accessed June 2026). If one sequence contained a matched on the +1 and +2 frame, it was considered and restored *aprD*, while those with just the +1 frame were considered as *aprD* pseudogenes. Bitscores of 50 were considered a significant match, and the top two hits at the species level were kept reducing graphing very closely related strains. To examine the gene neighborhoods of bacteria harboring a Psh protein, the GFF’s of bacteria with full genome assemblies were compiled, and a sliding search for annotated genes was conducted using the queries in **Supplemental Table 4**.

To find the cargo of genes that had a DUF4214 domain but lacked the RTX repeat transport found in the *P. putida* Psh protein, the amino acid sequences of those proteins were queried with Interpro (July 2026). The results were compiled and sorted with the following eight categories (listed as category and terms consolidated into that category): 1) DUF4214 Core Domain (C-term) (PF13946), 2) DUF-like / Phycobilisome Superfamily (G3DSA:1.10.3130.20), 3) Heme Peroxidase / Oxidase Domain (G3DSA:1.10.640.10, PF03098, PTHR11475, cd09821), 4) Serralysin-Like Metallopeptidase (G3DSA:3.40.390.10, SSF55486, SM00235, PF08548, PF00413, cd04277), 5) S-Layer Domain Signature (PTHR38340, IPR050557), 6) Cadherin Domain Repeat (Cadherin, PS50268, G3DSA:2.60.40.60, PS51854, PF16184, SSF49313, SM00112, cd11304), 7) Ig-Fold Structural Antenna (Immunoglobulin, Ig-like, G3DSA:2.60.40.10, G3DSA:2.60.40.2700, PF19077, PF17963), 8) RTX Calcium-Binding Repeats/Motifs (PF00353, PR00313, PS00330)

### In Silico Structural Modeling and Interaction

To characterize the molecular interactions and structural stability of the Psh protein complex, computational structure and affinity predictions were generated using the Boltz-2 structural biology foundation model (https://github.com/jwohlwend/boltz). The primary sequence of the Psh protein (NCBI protein ID AAN67414.1) was evaluated against modified lipopolysaccharide (LPS) components. The baseline core of the LPS ligand was constructed as a uniform 9- rhamnose carbohydrate chain (SMILES - C[C@H]1O[C@H](O[C@@H]2[C@@H](O)[C@H](O)O[C@@H](C)[C@@H]2O[C@@H]3[C@@H](O)[C@H](O)O[C@@H](C)[C@@H]3O[C@@H]4[C@@H](O)[C@H](O)O[C@@H](C)[C@@H]4O[C@@H]5[C@@H](O)[C@H](O)O[C@@H](C)[C@@H]5O[C@@H]6[C@@H](O)[C@H](O)O[C@@H](C)[C@@H]6O[C@@H]7[C@@H](O)[C@H](O)O[C@@H](C)[C@@H]7O[C@@H]8[C@@H](O)[C@H](O)O[C@@H](C)[C@@H]8O[C@@H]9[C@@H](O)[C@H](O)O[C@@H](C)[C@@H]9O)[C@ @H](O)[C@@H](O)[C@@H]1O).

The LPS ligand with a pseudaminic acid cap SMILES structure was CC(=O)N[C@@H]1[C@H](O[C@@](O[C@@H]2[C@@H](O)[C@H](O)O[C@@H](C)[C@@H]2O[C@@H]3[C@@H](O)[C@H](O)O[C@@H](C)[C@@H]3O[C@@H]4[C@@H](O)[C@H](O)O[C@@H](C)[C@@H]4O[C@@H]5[C@@H](O)[C@H](O)O[C@@H](C)[C@@H]5O[C@@H]6[C@@H](O)[C@H](O)O[C@@H](C)[C@@H]6O[C@@H]7[C@@H](O)[C@H](O)O[C@@H](C)[C@@H]7O[C@@H]8[C@@H](O)[C@H](O)O[C@@H](C)[C@@H]8O[C@@H]9[C@@H](O)[C@H](O)O[C@@H](C)[C@@H]9O[C@@H]10O[C@@H](C)[C@H](O)[C@H](O)[C@H]10O)(C[C@H]1NC(=O)C)C(=O)O)[C @@H]([C@@H](C)O)O.

To test whether Psh protein is predicted to bind to LPS, calcium, and the pseudaminic acid cap, the following conditions were tested: the absence or presence of localized divalent calcium Ca^2+^ ions, the absence or presence of a terminal pseudaminic acid moiety acting as the LPS structural cap. All structural assemblies were run using default parameters on the Boltz-2 architecture to derive 3D structural predictions and interface affinity calculations. The resulting chemical complexes were systematically quantified using three diagnostic scoring metrics: the predicted negative log half-maximal inhibitory concentration equivalent (PIC_50_), the Interface Predicted Distance Error (IPDE), and the Interface Predicted TM-score (IPTM). Structural visualizations and surface mapping fields were rendered from the highest-confidence PDB outputs for representative combinations.

### Statistical Analyses

To obtain a single luminescence value for each replicate across the kinetic luminescence curve for HiBit tagged proteins, the maximum value of the kinetic curve was taken from each replicate and condition. To calculate the secretion efficiency (Export Ratio), the mean total extracellular luminescence was divided by the mean cell lysate luminescence (adjusted by a factor of 10 to account for sample dilution). Standard error of the mean (SEM) for the resulting ratios was determined via standard relative error propagation of the individual sample variances (n = 3 replicates).

To provide a single representative sequencing metric for whole genome sequencing validation, a genome-wide weighted average coverage *C_w_* was computed across all assembled segments for each isolate, utilizing the following formula:

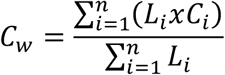

where *L_i_* represents the length of each contig (measured in base pairs), and *C_i_* is the depth of coverage for that contig. Final calculated weighted average coverages were integrated directly into the downstream comparative visualization matrices.

An unpaired two-sided student’s t-test was performed to compare transformation efficiencies between wild-type *P. putida* KT2440 and EmajogiR.1. Transformation efficiencies were calculated by dividing the CFU counts of transformed cultures plated on LB media versus the CFU counts of transformed cultures plated on LB plates with Kanamycin added.

## Acknowledgments

We thank Jesse Cahill for sharing the pJLC140 expression plasmid backbone used in this study, and Jeff Wang, Jessica Trinh, and Hazel Sisson for insightful discussions. We thank Lillian Lowrey for internal review of this manuscript. We thank the DOE Joint BioEnergy Institute and Alberto Rodriguez for providing the hydrolysates used in this work. We thank Hedvig Tamman for providing the CEPEST phage collection used in this work.

Research funding provided by Laboratory Directed Research and Development program at Sandia National Laboratories (233066 to CMM).

Sandia National Laboratories is a multimission laboratory managed and operated by National Technology & Engineering Solutions of Sandia, LLC, a wholly owned subsidiary of Honeywell International Inc., for the U.S. Department of Energy’s National Nuclear Security Administration under contract DE-NA0003525. This paper describes objective technical results and analysis.

Any subjective views or opinions that might be expressed in the paper do not necessarily represent the views of the U.S. Department of Energy or the United States Government. This article has been authored by an employee of National Technology & Engineering Solutions of Sandia, LLC under Contract No. DE-NA0003525 with the U.S. Department of Energy (DOE). The employee owns all right, title and interest in and to the article and is solely responsible for its contents. The United States Government retains and the publisher, by accepting the article for publication, acknowledges that the United States Government retains a non-exclusive, paid-up, irrevocable, world-wide license to publish or reproduce the published form of this article or allow others to do so, for United States Government purposes. The DOE will provide public access to these results of federally sponsored research in accordance with the DOE Public Access Plan https://www.energy.gov/downloads/doe-public-access-plan.

## Author Contributions

B.P.D. and C.M.M. conceived the experiments and designed experiments. B.P.D., J.D.J, S.W. performed research. M.R. and V.R.C provided novel reagents for the experiments. B.P.D. performed all statistical analysis and developed figures. C.M.M. supervised the research and obtained the funding. B.P.D, S.W., J.D.J, M.R wrote the manuscript. All author authors revised and edited the manuscript.

