## Supplemental Figure for "Phage proofing *Pseudomonas putida* uncovers novel broad-spectrum phage resistance protein Psh"

**Supplemental Information**

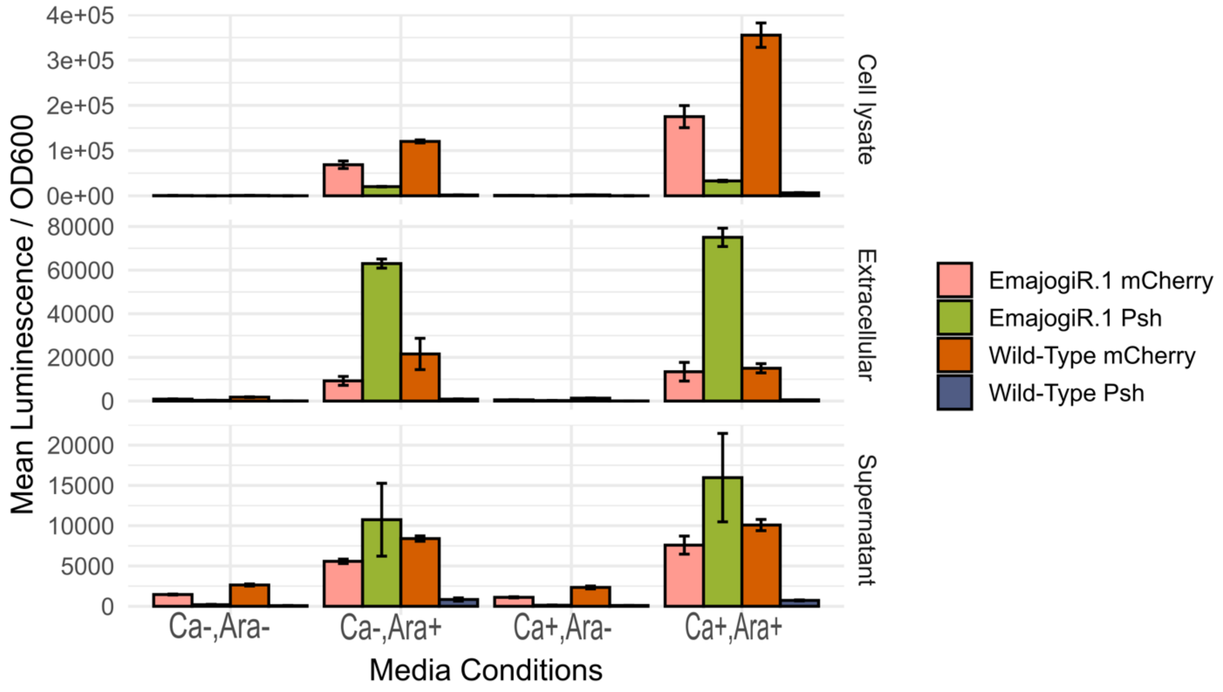


**Supplemental Figure 1. Non-normalized fractional HiBit-tagged Psh protein expressed in wild-type *P. putida* and EmajogiR.1 strains.** Error bars represent the SEM of each set of values. Proteins were expressed on an arabinose-inducible construct: "Arabinose +" refers to 0.2% arabinose expression for 3.5 hours before this assay. Ca+ is the addition of 5mM $CaCl_2$ and Ara+ is the addition of 0.2 % arabinose.

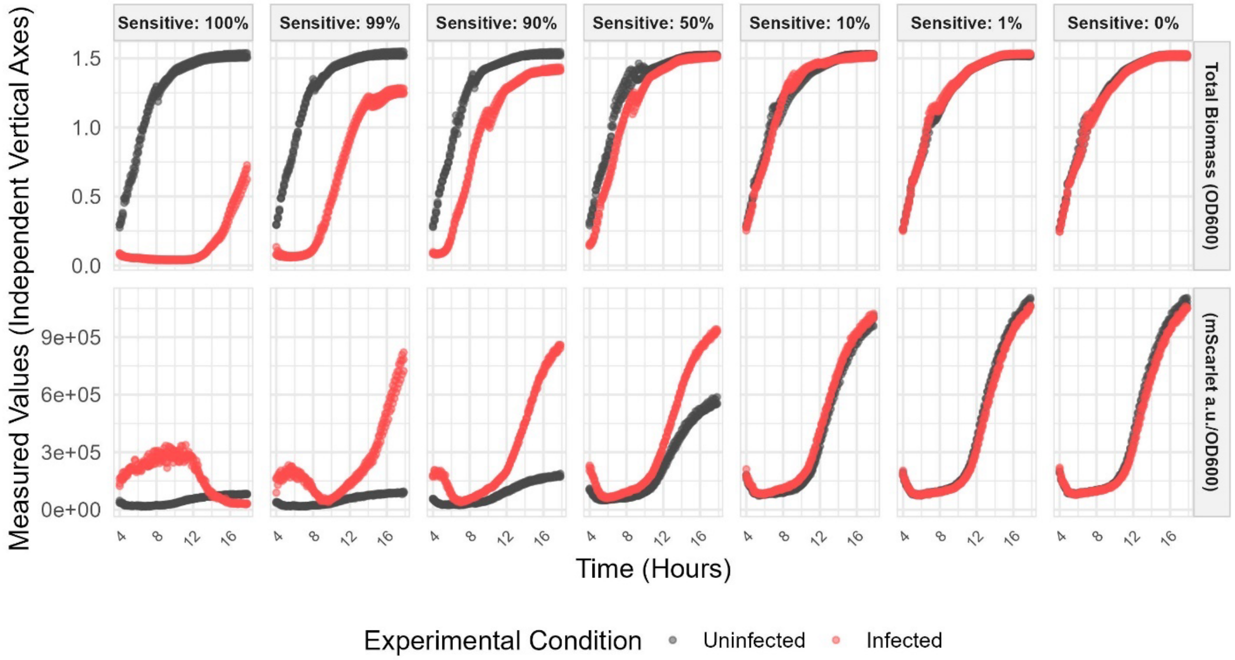


**Supplemental Figure 2. Coculture of phage-resistant mutant does not protect infection sensitive strain.** Wild-type *P. putida* KT 2440 and *EmajogiR.1* expressing mScarlet were combined at different rations and infected with EmajogiR.1 at an MOI of 0.001. The ratio of mScarlet to OD600 represents the fraction of resistant EmajogiR.1 that's present. Note that in the left most panel, some background fluorescence shows low levels of mScarlet/OD600. The time listed is hours post infection with Emajogi. Red denotes infected cultures and black denotes uninfected cultures.

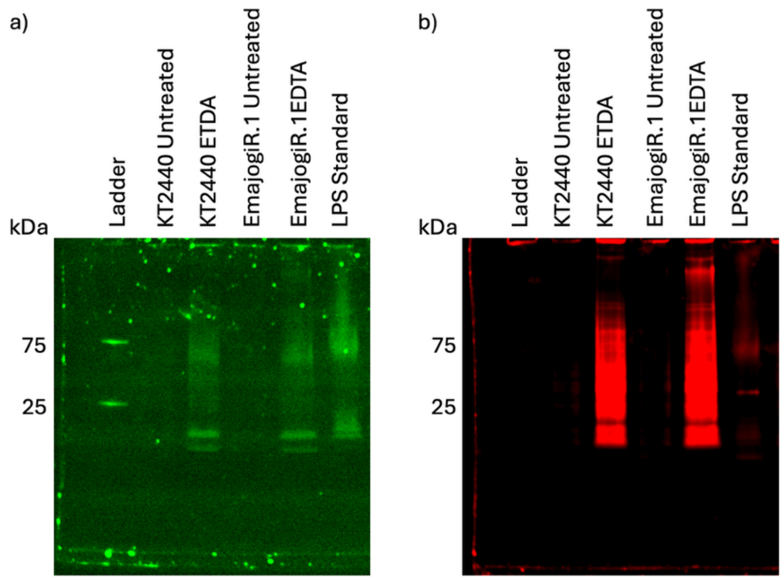


**Supplemental Figure 3. LPS and protein co-staining reveal large protein co-eluting in EmajogiR.1 LPS extractions.** SDS-PAGE gels with either *P. putida* KT2440 or *P. putida* EmajogiR.1 lysates. One aliquot was treated with 5mM EDTA to disrupt the LPS stained with **a)** Pro-Q Emerald 300 for total protein or **b)** SpyroRuby for LPS. Ladder is Precision Plus Protein Dual Color Standards and the LPS standard is ProQ (*E. coli* serotype 055:B5) 1:100 dilution (~2.5 µg).

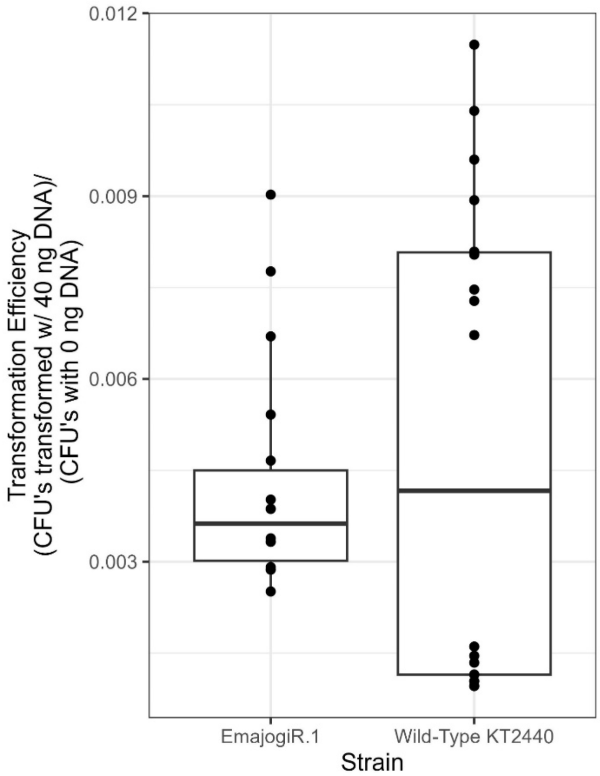


**Supplemental Figure 4. Transformation efficiency is the same between EmajogiR.1 and wild-type KT2440.** Transformation efficiency compared by transforming 40ng of pJLCKan/mCherry into both wild-type *P. putida* KT2440 and strain EmajogiR.1. Total CFU counts plated on LB agar supplemented with kanamycin were normalized to total cells transformed (CFU's with 0 ng DNA, plated on LB plates without antibiotics). A two-sided student's t-test was performed.

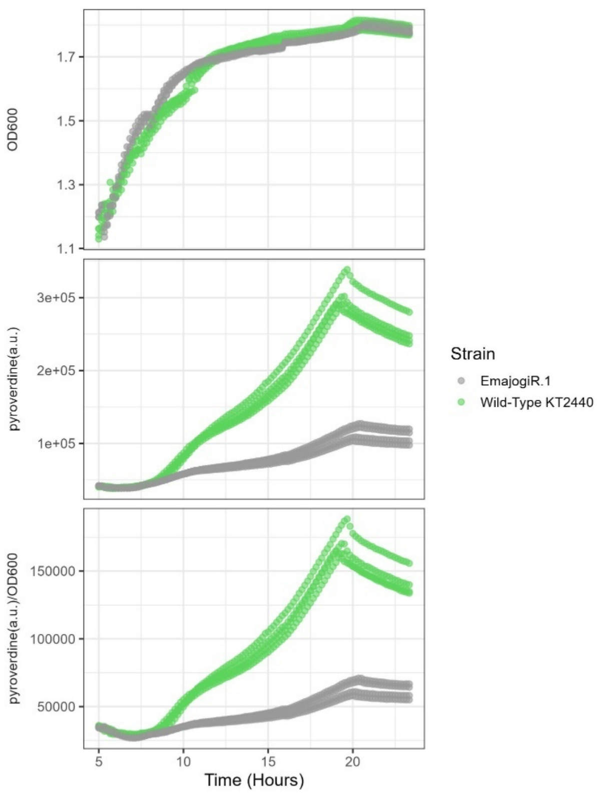


**Supplemental Figure 5. Pyoverdine production per cell is lower in EmajogiR.1 than Wild-Type *P. putida* KT2440.** Growth curves were taken from subcultured overnight cultures of EmajogiR.1 and wild-type *P. putida* KT2440 at the same time on a 96-well plate. Pyoverdine was measured as Ex. 400(± 10), Em. 460 (± 15). (n=3)

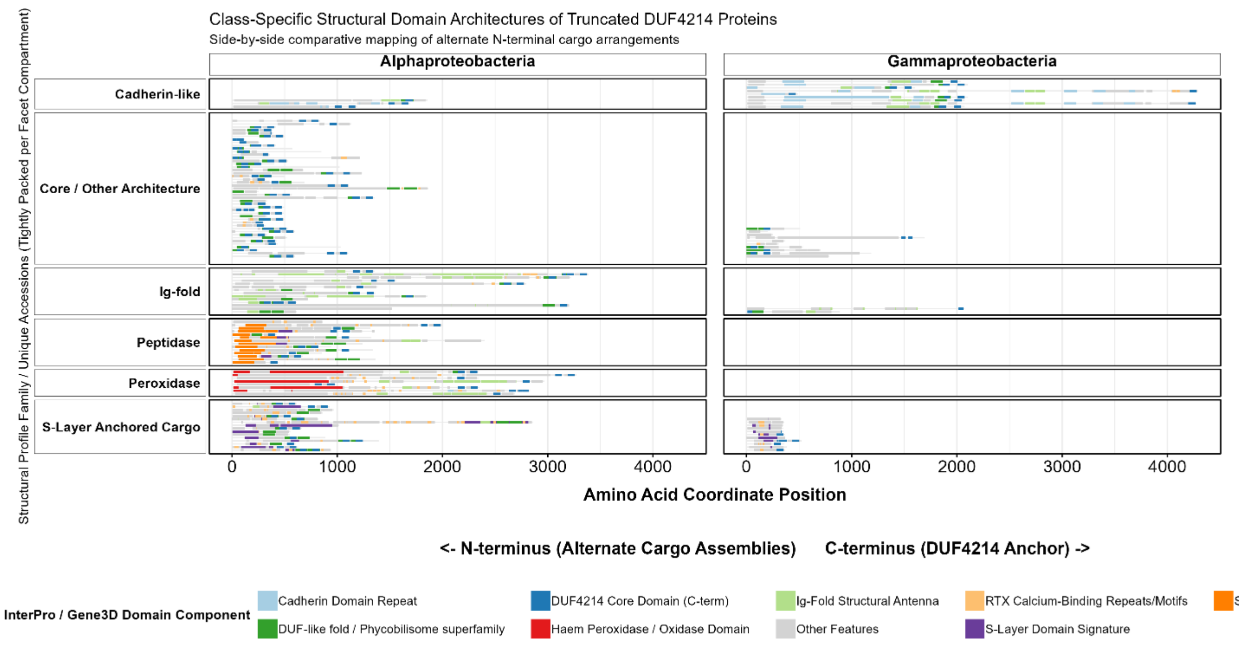


**Supplemental Figure 6. Class-Specific structural domain architectures of proteins containing DUF4214 domains in *Alpha*- and *Gammaproteobacteria*.** Proteins only containing the DUF4214 domain were mapped to discover other domains associated with this domain. Alignments are displayed by amino acid coordinate position from the N-terminus to the conserved DUF4214 core domain at the C-terminus. Individual horizontal lines represent independent protein accessions. Faceted regions represent structural profile families classified by conservative domain combinations from the InterPro database. Panels compare biological distribution across the taxonomic classes *Alphaproteobacteria* (left) and *Gammaproteobacteria* (right), highlighting an asymmetry in N-terminus contents between lineages. Colored internal segments mark independent diagnostic feature coordinates.

a)

| $Ca^{2+}$ ions | LPS Cap | Binding affinity (PIC50) | Complex IPDE | Ligand IPTM |
|---|---|---|---|---|
| - | - | -9.8 | 8.9 | 0.45 |
| + | - | -9.8 | 7.9 | 0.52 |
| - | + | -9.8 | 8.5 | 0.51 |
| **+** | **+** | **-10.2** | **7.3** | **0.55** |

b)

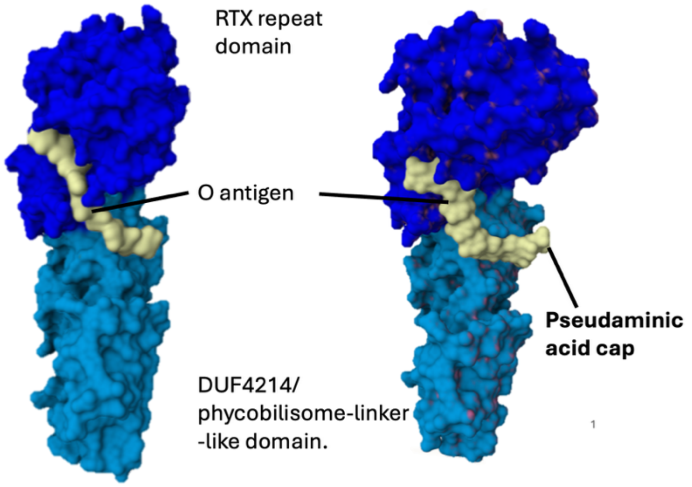


**Supplemental Figure 7. Structural modeling and binding affinity predictions of the Psh protein complex with lipopolysaccharide and divalent calcium. a)** Performance matrix displaying predicted binding affinity (pIC50), Interface Predicted Distance Error (IPDE), and Interface Predicted TM-score (IPTM) across four experimental states evaluated with or without divalent calcium (Ca2+) ions and the pseudaminic acid lipopolysaccharide (LPS) cap. Bold text highlights optimal stabilization metrics when both modifiers are present. **b)** 3D surface representations of the Psh protein complex derived from the structural modeling workflow. The structural domains are colored and annotated: RTX repeat domain (dark blue), O-antigen 9-rhamnose chain (light yellow), DUF4214/phycobilisome-linker-like domain (cyan), and the pseudaminic acid cap modification (pale green).

### Supplemental Tables

Supplementary Table 1. Minimum inhibitory concentrations of EmajogiR.1 and wild type across diverse antibiotics.

Supplementary Table 2. Primers used in this study.

Supplementary Table 3. Plasmids generated for this study.

Supplemental Table 4. Queries for gene neighborhood of Psh gene.
